# Keratin 5 marks cancer-propagating cells sustained by an osteopontin-producing niche in high-grade serous ovarian carcinoma

**DOI:** 10.64898/2026.01.28.702332

**Authors:** Mallikarjun Bidarimath, Coulter Q. Ralston, Nandini Bidarimath, Ian M. Rose, Darianna Colina, Elisa Schmoeckel, Andrew K. Godwin, Doris Mayr, Lora H. Ellenson, Andrea Flesken-Nikitin, Alexander Yu. Nikitin

**Author notes:** Correspondence to Andrea Flesken-Nikitin; or Alexander Nikitin.

## Abstract

High-grade serous carcinoma (HGSC) is the most common and aggressive form of ovarian cancer. Advanced HGSCs exhibit pronounced cellular heterogeneity, including a subset of cancer-propagating cells (CPCs, also known as cancer stem cells) that are highly tumorigenic and display stem cell-associated properties such as self-renewal and chemoresistance. In contrast, a substantial fraction of HGSC cells is non-tumorigenic. The role of these non-cancer-propagating cells (non-CPCs) and their relationship to CPCs remain poorly understood. Here, we demonstrate that neoplastic cells expressing the intermediate filament protein keratin 5 (KRT5) represent bona fide CPCs. KRT5⁺ cells form cancer organoids over successive passages, are tumorigenic in serial dilution xenograft assays, and are resistant to the antineoplastic agents, doxorubicin and cisplatin. Single-cell lineage-tracing experiments show that KRT5⁺ CPCs give rise to KRT5⁻ cells. KRT5⁺ and KRT5⁻ populations exhibit distinct gene expression profiles, with KRT5⁻ cells characterized by expression of *SPP1*, which encodes the secreted factor osteopontin (OPN). Treatment with OPN enhances HGSC organoid growth and chemoresistance, whereas *SPP1* knockdown reverses these effects. Together, these findings support a model in which HGSC contains two hierarchically related cell populations: KRT5⁺, OPN-responsive CPCs and KRT5⁻, non-tumorigenic cells that form a niche producing OPN. Targeting pathways that sustain both stem-like tumor cells and their supportive niche may enable reduced dosing of highly toxic chemotherapeutic agents while enhancing therapeutic efficacy in HGSC.

## Introduction

Ovarian/extra-uterine high-grade serous carcinoma (HGSC) is the most common and aggressive type of ovarian cancer [1, 2]. It often has no symptoms at early stages and over 80% of patients are diagnosed at advanced, usually incurable, cancer stages, when the tumors have already metastasized [3, 4]. The overall 5-year survival rate of patients with advanced epithelial ovarian cancer is 32% [5]. Standard chemotherapy can substantially reduce the size of the tumor and transiently improve patient’s health status. However, due to extensive cellular heterogeneity, a fraction of cancer cells evades chemotherapy and advance the disease progression after a brief period of dormancy [6–8].

The evolution of cancer cells and mechanisms responsible for the development of chemoresistance remain insufficiently elucidated. Some cancer cells are highly tumorigenic and called cancer propagating cells (CPC), cancer stem cells or cancer-initiating cells [2, 9–12]. These cells feature stem cell-associated properties, such as self-renewal and chemoresistance. Several CPC markers, such as CD44, CD24 and CD133, have been reported [13–15]. However, their prognostic value and stability of associated phenotypes remain debatable [11, 16–18].

At the same time, a significant fraction, in many cases the majority, of cancer cells has low, if any, tumorigenicity. The role of such non-cancer propagating cells (non-CPC) in HGSC progression and chemoresistance, and cellular dynamics between CPC and non-CPC remain largely unknown. Non-CPC can be considered as a byproduct of CPCs’ stem cell-like ability to differentiate. However, non-CPC may also have niche-like properties typical for the normal tissue homeostasis [9, 19–21].

Previously it has been reported that epithelial ovarian cancer cell lines expressing the intermediate filament protein keratin 5 (KRT5) are more resistant to cisplatin-induced apoptosis [22]. Furthermore, it has been suggested that KRT5 overexpression is associated with serous ovarian cancer recurrence and chemotherapy resistance [23]. However, it remains unknown if KRT5+ cells represent CPC.

Our studies found that KRT5+ cells show features typical for CPC, such as high organoid forming capacity in consecutive passages, tumorigenicity in serial dilutions, ability to generate KRT5- progeny, and chemoresistance. Notably, growth and chemoresistance of KRT5+ CPC is sustained by non-CPC-produced osteopontin (OPN), a secreted, sialic acid-rich, glycosylated phosphoprotein encoded by *SPP1* gene [24].

## Results

### Clinical relevance of KRT5+ cells in HGSC

Previous studies reported presence of KRT5+ cells in OV432, SKOV3, OVCAR3, PEO1, and PEO4 cell lines at frequencies between 2.4% to 52.7% [22]. Consistent with this observation we observed a subpopulation of KRT5+ cells in cell lines CAOV-3 (1.82%), CAOV-4 (4.99%) and SKOV3 (7.91%) (Table S1).

In Kaplan-Meier Plotter *KRT5* mRNA expression was associated with a poor prognosis of patients with serous ovarian carcinoma with TP53 mutation (Figure S1). According to scRNA seq datasets, all HGSC (n=29) contained KRT5+ cells (Table S2) and 48% (14 out of 29) HGSC contained at least 5% of KRT5+ cells. We further confirmed KRT5 expression by immunostaining in 197 HGSC cases with known clinical outcomes (Figure 1). Higher frequency of KRT5+ cells (≥40% of labeled cells) was associated with significantly shorter survival period after diagnosis (KRT5+ vs KRT5-, 955 days vs 1452 days, Logrank P=0.0015). Overall, these observations are consistent with the previous report based on 117 patients [23].

**Figure 1.**
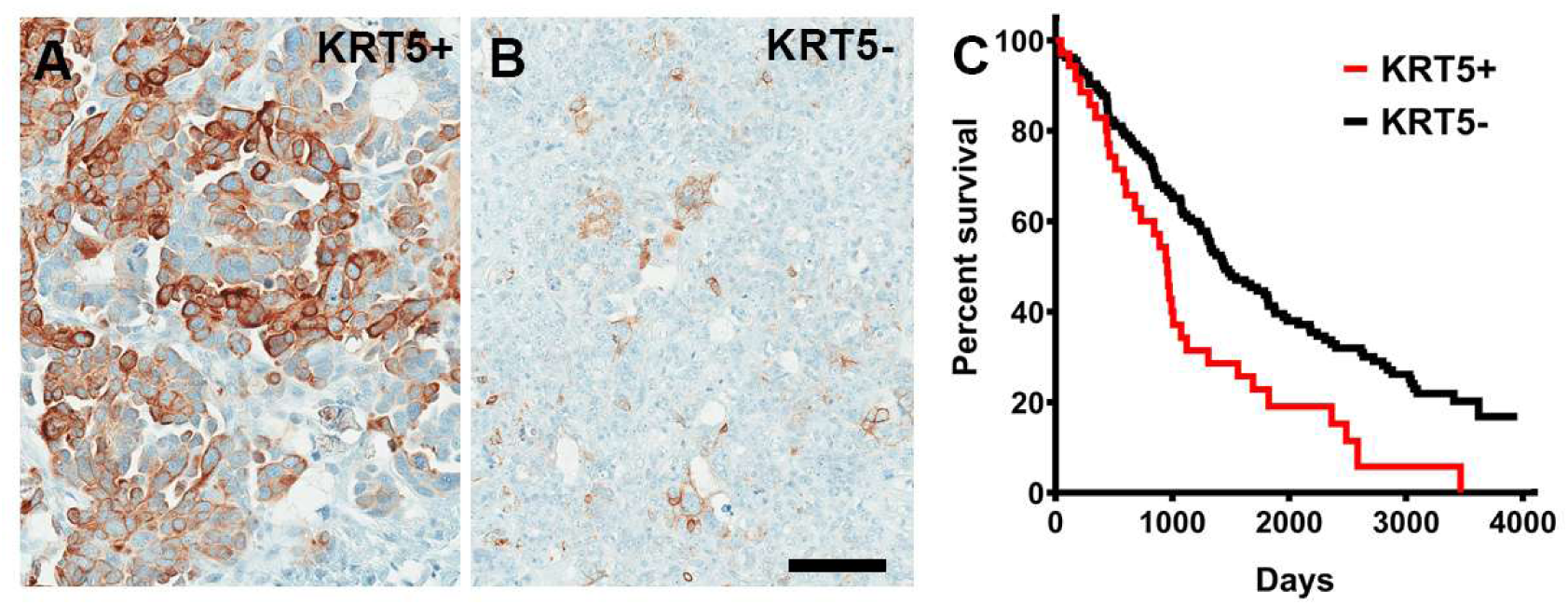
KRT5+ cells in HGSC. **A** and **B**, High (A, KRT5+, >40% stained neoplastic cells, n = 37) and low (B, KRT5-, n = 160) frequency of KRT5 expressing cancer cells. **C**, Kaplan-Meier survival analysis of HGSC patients stratified according to frequency of KRT5 expressing cells (P=0.0015). A and B, Elite ABC method. Hematoxylin counterstaining. Scale bar, 100 µm.

### Isolation and characterization of KRT5+ cells

To gain insights into the biological properties of KRT5+ cells we transduced SKOV3 cells with lentiviruses containing *KRT5* promoter driving expression of reporter mCherry (Figure 2A-C). Over 95% of KRT5+ cells expressed mCherry (Figure 2B and C). No expression of mCherry was observed in KRT5- cells. To aid in identification of KRT5+ cell progenies irrespective of whether KRT5+ expression has been retained in daughter cells, cells were co-transduced with lentivirus expressing GFP driven by the ubiquitous *hUbC* promoter (L-hUbC-GFP, Figure 2D). We then isolated KRT5+ and KRT5- cell populations using fluorescence activated cell sorting (FACS). According to PCR analysis, both lentiviruses evenly integrated in both KRT5+ and KRT5- cells. (Figure 2E). Thus, our approach provides a platform to test both KRT5+ and KRT5- cell populations for their functional properties.

**Figure 2.**
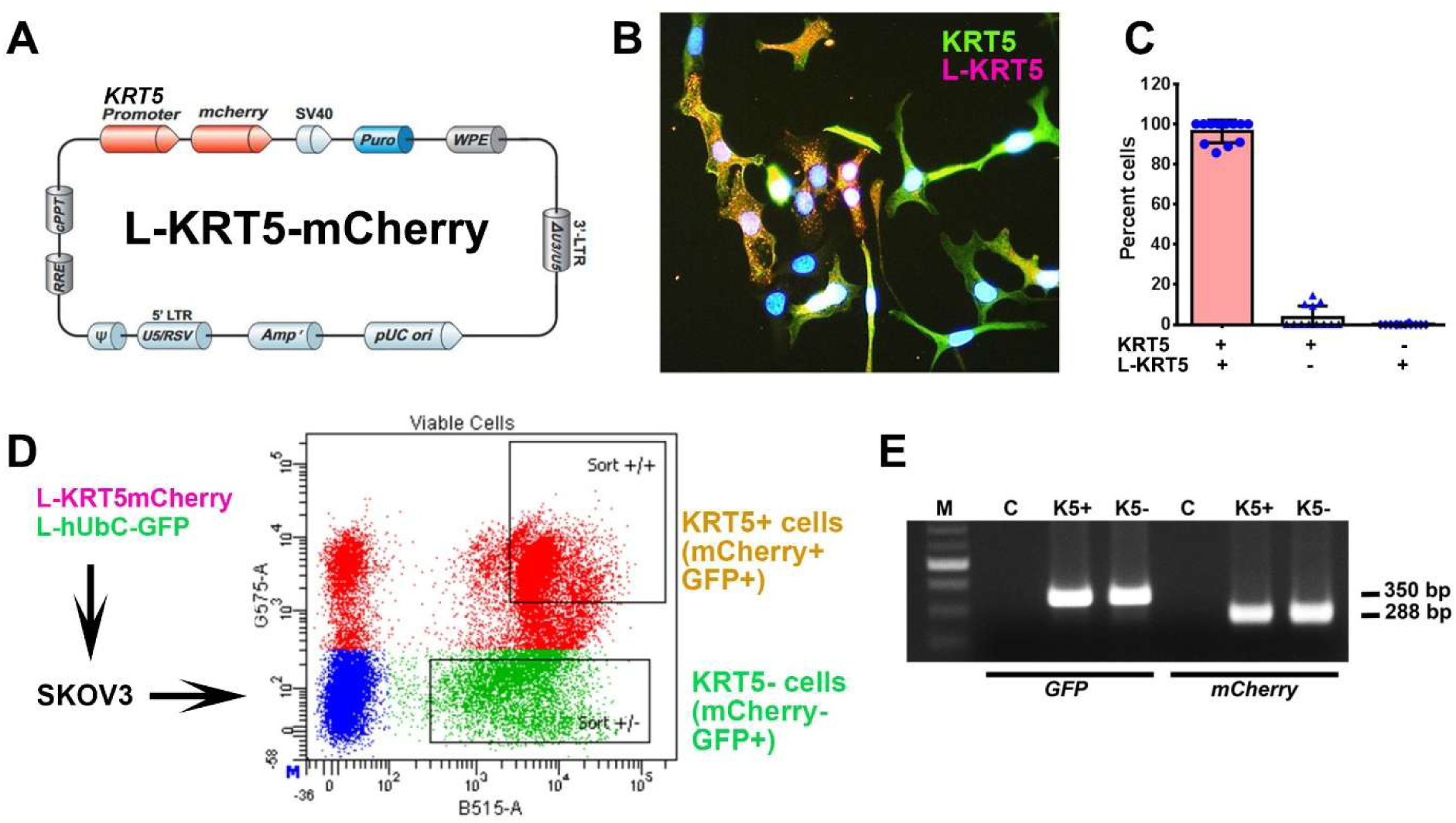
Isolation and characterization of KRT5+ and KRT5- cells from SKOV3 cell line. **A**, Structure of Lenti-KRT5mCherry (L-KRT5mCherry). **B**, Co-localization of KRT5 immunostaining (KRT5, green) and mCherry (L-KRT5) expression (magenta). Orange, overlay. Counterstaining with DAPI (blue). Scale bar, 60 µm. **C**, Quantification of cells expressing both KRT5 and L-KRT5 (KRT5+ L-KRT5+), or only KRT5 (KRT5+) or L-KRT5+. **D**, Experimental design of isolation of cells expressing both L-KRT5mCherry and L-hUbC-GFP or L-hUbC-GFP alone. **E,** PCR detection of L-hUbC-GFP (GFP) and L-KRT5mCherry (mCherry) DNA in both KRT5+ (K5+) and KRT5- (K5-) cells. All error bars denote s.d.

KRT5+ cells successfully generated cancer organoids for 6 passages, with each passage lasting 12 days, following seeding in Matrigel. In contrast, KRT5- cells were only capable of forming organoids for up to 3 passages (Figure 3A-C). Compared with KRT5-cells, organoids derived from KRT5+ cells were significantly larger (Figure 3A and B) and formed at a higher frequency (Figure 3C). Similar results were obtained after growing organoids in ultra low attachment plates (Figure S2).

**Figure 3.**
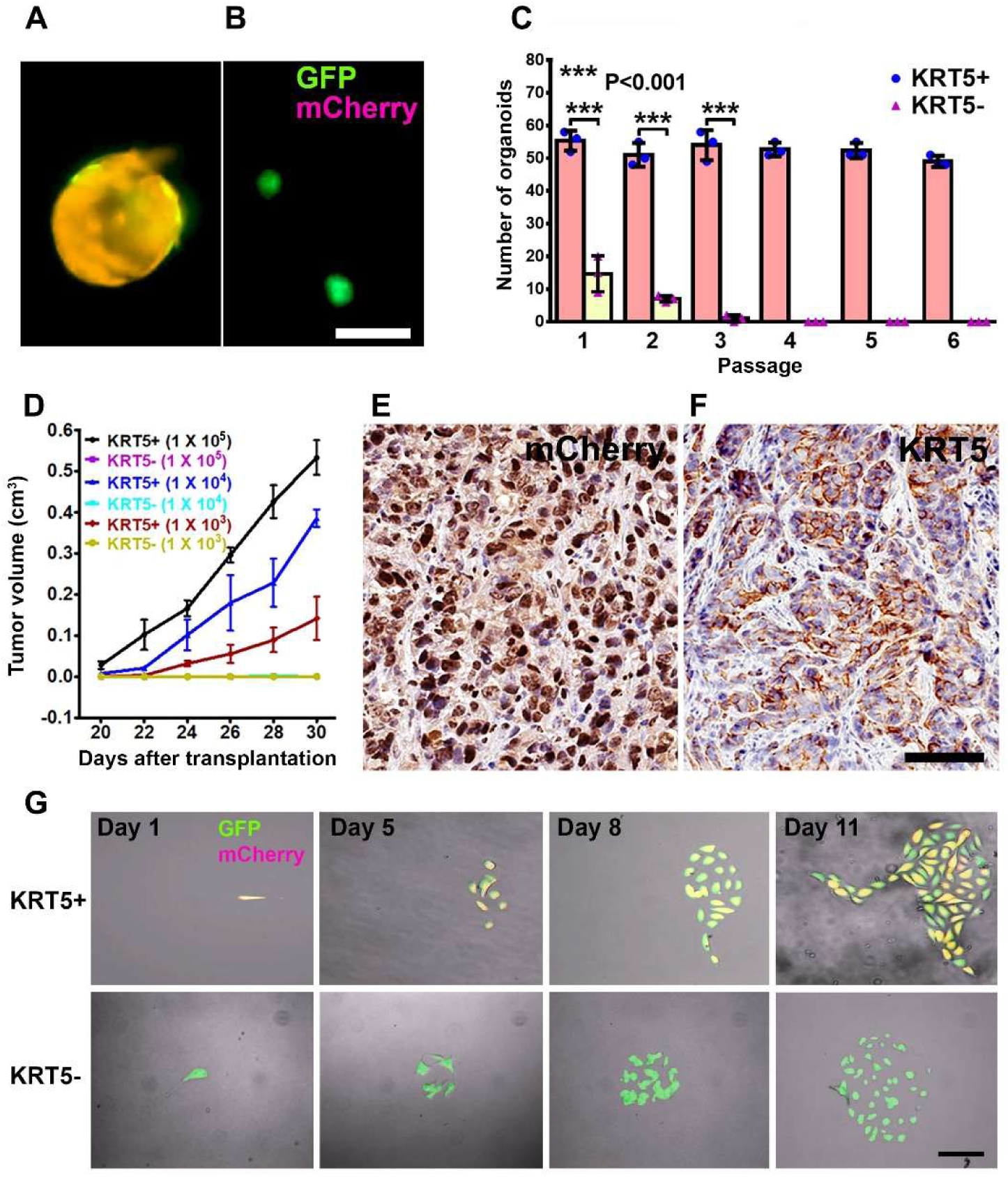
Characterization of KRT5+ cells. **A** and **B**, Organoids derived from SKOV3 cells expressing both lentiviruses (A, GFP and mCherry, orange) or GFP alone (B, green). Scale bar, 100 µm. **C**, Quantification of KRT5+ (blue symbols, pink bars) and KRT5- (pink symbols, yellow bars) cancer organoids in 6 consecutive passages. All error bars denote s.d. **D,** Volume of tumors formed by serially diluted (1 x 10^5^, 1 x 10^4^, 1 x 10^3^) of KRT5+ and KRT5- cells after their s.c. transplantation into different flanks of NSG mice. KRT5-group did not form tumors. **E** and **F**, mCherry (E) and KRT5 (F) expression in KRT5+ cell derived xenografts. Elite ABC method. Hematoxylin counterstaining. Scale bar, 60 µm. All error bars denote s.d. **G.** Live microscopy of cells were isolated by FACS based on their expression of GFP (green) and mCherry (magenta) after coinfection with Lenti-UbC-GFP and Lenti-KRT5mCherry. Orange, Overlay. Individual frames of live microscopy. Scale bar, 60 µm.

To assess the tumorigenicity potential of KRT5+ and KRT5- cells in vivo, FACS-isolated and serially diluted (1 x 10^5^, 1 x 10^4^, 1x 10^3^) KRT5+ and KRT5- cells were subcutaneously transplanted into NOD scid gamma (NSG) mice. Beginning 20 days post-transplantation KRT5+ cells formed tumors in a dose dependent manner, whereas KRT5-cells failed to form tumors by 30 days after transplantation (Figure 3D; Table S3). Tumor tissues were collected at the experimental endpoint on day 30 days and subsequently validated for mCherry and KRT5 expression by immunostaining (Figure 3E and F).

To test if KRT5+ cells give rise to KRT5- cells and vice versa, we seeded single KRT5+ or KRT5- cells in a 35 mm µ-Dish with a glass bottom and followed their fate at 24 hours intervals for 8-10 days using a fluorescent microscope (Figure 3G). KRT5+ cells gave rise to KRT5- cells (n=3 independent tracing experiments). However, KRT5- cells did not revert into KRT5+ cells (n=3 independent tracing experiments).

### KRT5+ cells have increased resistance to anti-cancer drugs

To compare chemoresistance of KRT5+ and KRT5- cells they were treated with increasing concentrations (2.5 µM, 5.0 µM and 10.0 µM) of common antineoplastic drugs cisplatin and doxorubicin (Figure 4A). According to both manual counting and MTT assay, there was a significant decrease in the cell viability of both KRT5+ and KRT5- cells in a dose dependent manner 72 hours after treatment. However, KRT5+ cells were significantly more resistant, with approximately 35%-40% viable cells after treatment with the highest concentrations of either cisplatin or doxorubicin.

**Figure 4.**
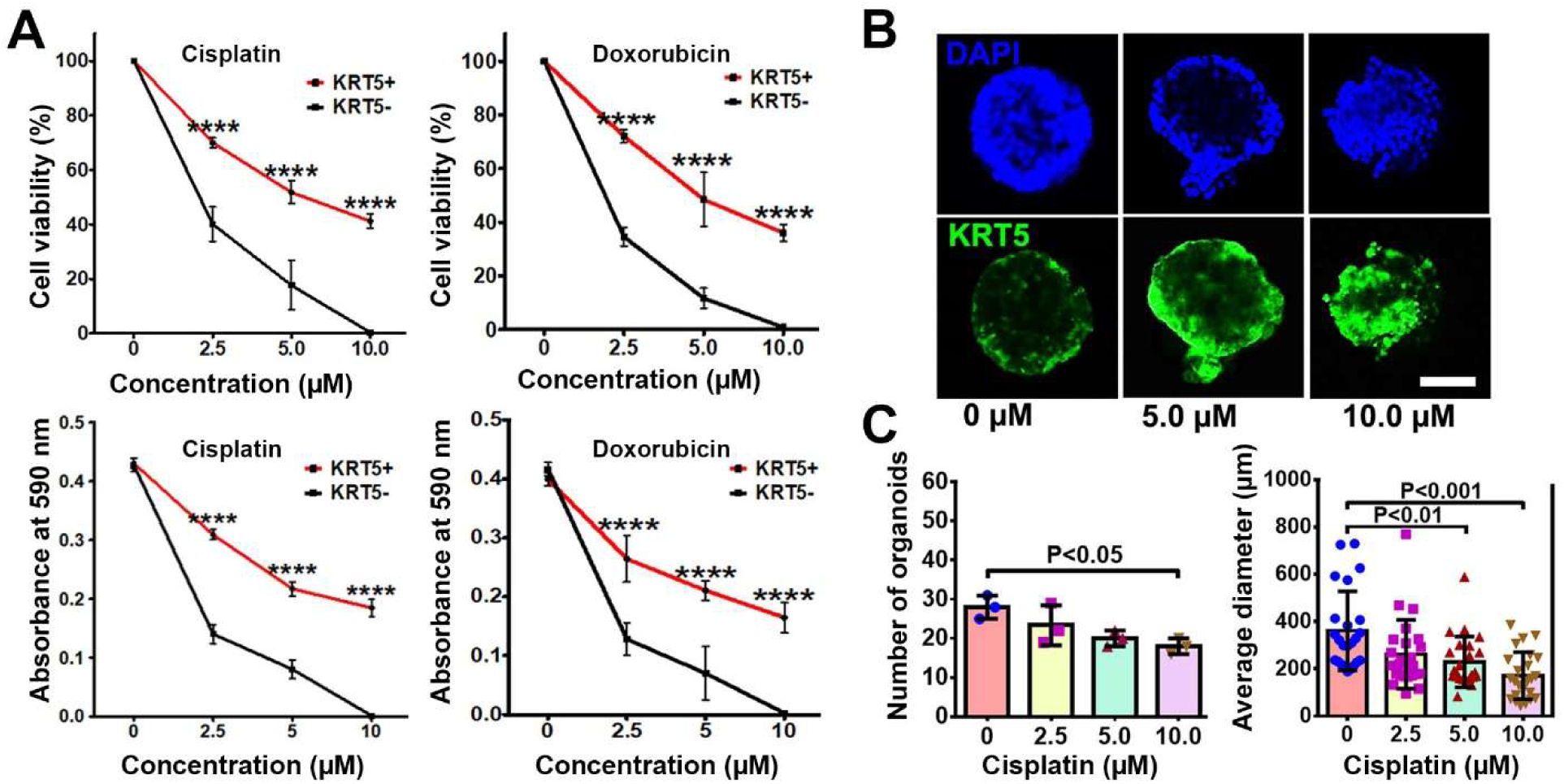
KRT5+ cells are chemoresistant to anti-cancer drugs. **A.** Effect of cisplatin and doxorubicin on KRT5+ and KRT5- cells. Manual count (cell viability, upper row) and MTT assay (absorbance at 590 nm, bottom row) of KRT5+ and KRT5- cells after 72 hours of treating with cisplatin and doxorubicin at different concentrations. **B** and **C**. Representative confocal images of primary HGSC organoids (B) and quantification of organoid frequency (B, left image) and size (C, right image) after treatment with different concentrations of cisplatin for 72 hours. C, Immunofluorescence for KRT5 (green). Counterstaining with DAPI (blue). Scale bar, 50 µm for all images. All error bars denote s.d.

To confirm the translational relevance of our findings, we evaluated effect of drugs on KRT5 expression in three KRT5+ tumorigenic organoid systems established by us from individual HGSC samples. In agreement with observations on cancer cell lines, treatment of cancer organoids with increasing concentrations of cisplatin resulted in reduction of organoid numbers and their sizes (Figure 4B and C). At the same time, the relative number of KRT5+ cells increased.

### Distinct gene expression signatures of KRT5+ and KRT5- cells

To compare transcriptomes of KRT5+ and KRT5- cells, we sorted 3 independent samples by FACS followed by RNA sequencing and visualized the top 200 differentially expressed genes distinguishing KRT5+ and KRT5- groups (Figure 5A). According to DAVID bioinformatics analysis, KRT5+ cells’ transcriptomes were significant in transcription regulation, cell division/cycle, cell-cell adhesion, and DNA damage and repair annotations (Figure 5B). Functional GO enrichment of KRT5- cells reveal that the top significant terms were observed in secreted, signal peptide, and extracellular-related annotations (Figure 5C).

**Figure 5.**
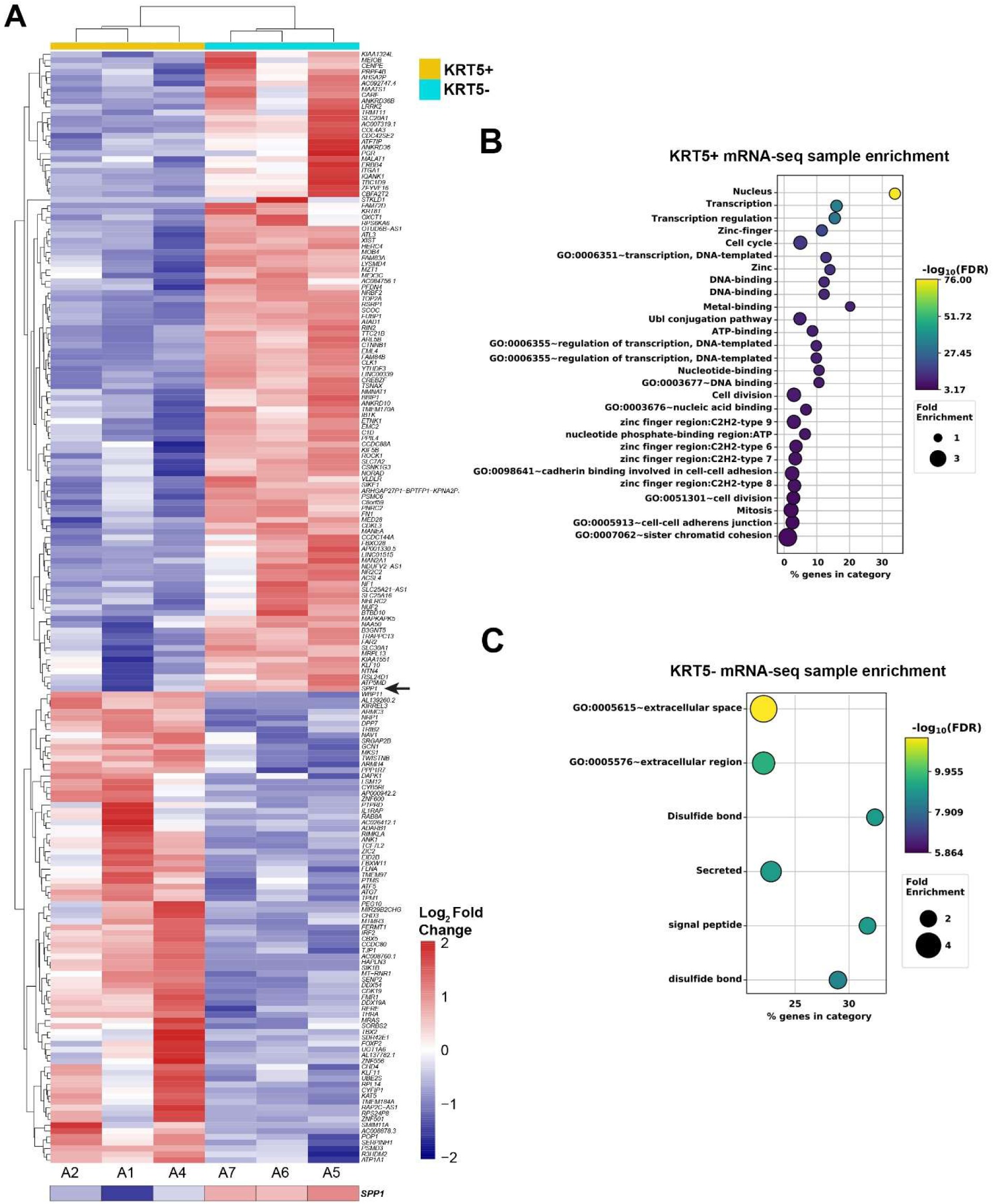
Gene expression analysis of KRT5+ and KRT5- cells isolated from SKOV3 cell line. **A**. Heat map depicting the top 200 differentially expressed genes in KRT5+ (A2, A1 and A4) and KRT5- (A7, A6, and A5) cells. Note the expression of *SPP1* (osteopontin) in KRT5- cells (arrow). **B** and **C**. DAVID analysis for functional annotation of terms associated with KRT5+ (B) and KRT5- (C) cells.

### KRT5- cells produce OPN facilitating KRT5+ CPC maintenance and chemoresistance

During RNA sequencing analysis we identified high expression levels of *SPP1*, the gene encoding osteopontin (OPN), in KRT5- cells (Figure 5A) These data were corroborated with publically available single-cell RNA sequencing data from 41 patients with HGSC. Within these datasets, *KRT5* and *SPP1* co-expression rarely was reported, and a majority of *KRT5*+ epithelial cancer cells did not express *SPP1* and vice versa (Figure 6A and B).

**Figure 6.**
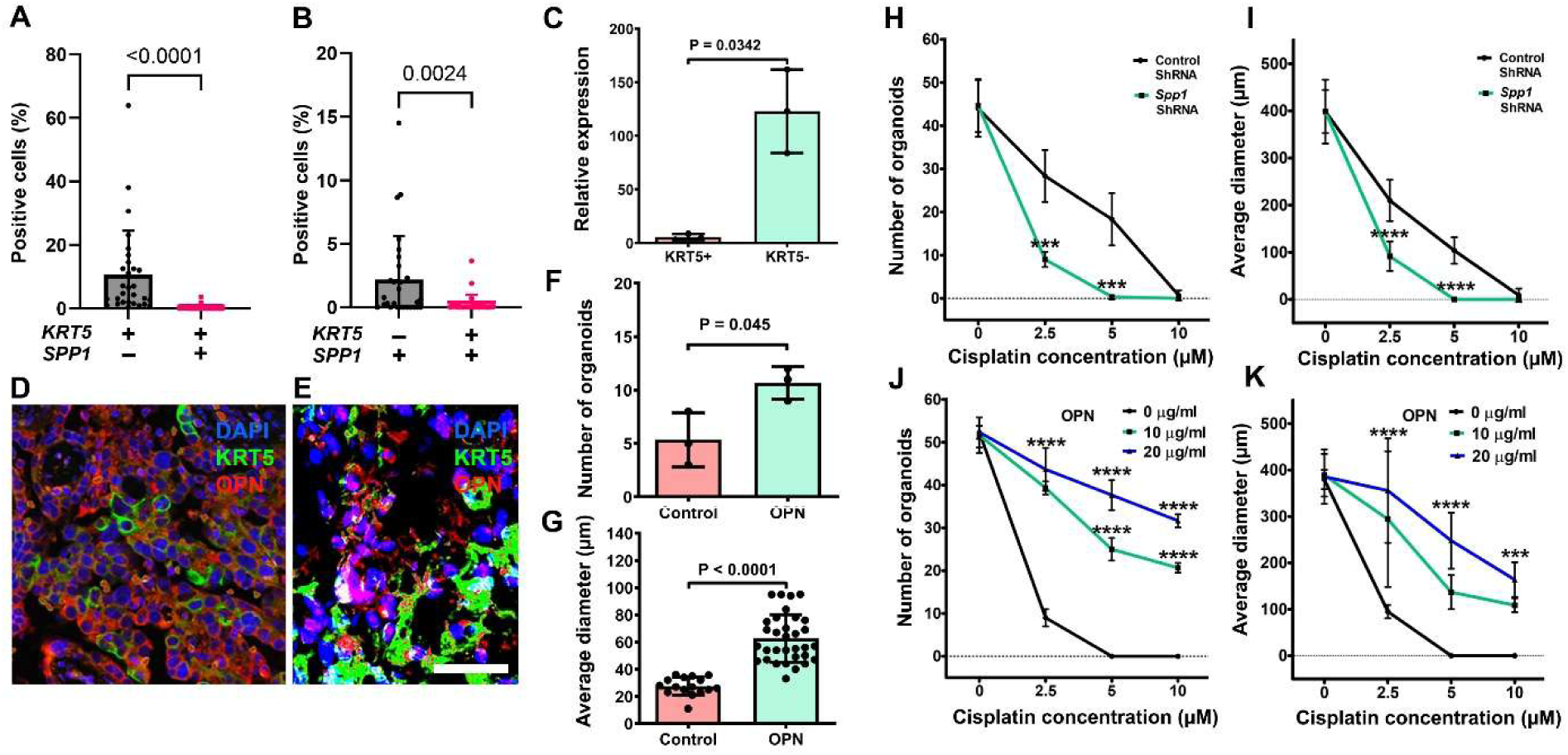
Characterization of OPN role in HGSC. **A** and **B.** Quantification of cells expressing *KRT5* and/or SPP1 cells within human HGSC cases from single-cell RNA sequencing. Significance by Mann-Whitney U test. **C.** RT-PCR analysis of *SPP1* expression in KRT5+ and KRT5- subpopulations of SKOV3 cells. **D** and **E,** KRT5 (green) and OPN (red) expression in HGSC (D) and primary HGSC organoid (E). Double immunofluorescence, counterstaining with DAPI (blue). Scale bar, (D) 60 µm, E (40 µm). **F** and **G**, OPN treatment increases frequency (F) and size (G) of HGSC organoids (n=3). **H** - **K**. Effect of cisplatin on frequency (H and J) and size (I and K) of organoids either transduced with *SPP1* shRNA (H and I) or treated with OPN (I and K). All organoids were measured 72 hours after treating with cisplatin at different concentrations. All error bars denote s.d.

Preferential *SPP1* expression in KRT5- cells was confirmed by RT-PCR of sorted cells (Figure 6C) and by immunostaining of primary HGSC (Figure 6D, n=8) and HGSC organoids (n=3) derived from the same cases (Figure 6E). Supporting biological significance of these observations, addition of OPN increased HGSC organoid frequency and size (Figure 6F and G). Importantly, *SPP1* knockdown by shRNA potentiated effects of cisplatin on frequency and size of organoids (Figure 6H and I), whereas increased OPN concentration had opposite effects in a dose-dependent manner (Figure 6J and K).

## Discussion

Heterogeneity in cancer cellular states is closely associated with the propagation of CPC. Here, we identify KRT5 expression as a reliable marker of CPC in HGSC. This observation is consistent with previous reports [22, 23] and our current findings that HGSC tumors with a high proportion of KRT5-positive cells exhibit more rapid progression and increased resistance to chemotherapeutic agents.

KRT5 is a well-established marker of stem/progenitor cells in lung and prostate epithelia [25, 26]. Its expression has been also detected in fallopian epithelial cells with stem-like properties, including the expression of epithelial cell adhesion molecule (EpCAM), CD44, and integrin α6, and the capacity for clonal growth and self-renewal in sphere formation assay [27]. To date, KRT5 expression has not been reported in serous tubal intraepithelial carcinoma. Notably, in mouse models *Krt5* expression marks pre-ciliated tubal epithelial cells that are prone to HGSC initiation [28]. These findings raise important questions regarding the evolutionary relationship between normal and neoplastic KRT5+ cell populations, which remain to be elucidated.

Hierarchies of cellular differentiation and the associated interrelationship among the tumor cells may determine the overall growth of the tumors and their susceptibility to drugs and targeted therapeutic approaches [19, 29–33]. Our study shows that HGSC contains two hierarchically related cell populations: KRT5⁺ CPCs and KRT5⁻ non-CPC cells. These cell populations have distinct transcriptome profiles that should allow in-depth evaluation of pathways associated with either tumorigenicity CPC or niche-producing factors by non-CPC.

Non-CPC cells produce OPN and thereby support growth and chemoresistance of CPC by activating OPN signaling pathway. Osteopontin (OPN), a secreted, sialic acid-rich, glycosylated phosphoprotein encoded by *SPP1* gene [24]. OPN interacts with cell surface receptors, such as CD44 and integrins [34, 35], and plays an important role in a wide range of physiological and pathological processes, such as cell adhesion, proliferation, survival, and angiogenesis. OPN signaling is known to regulate the stem/progenitor cell proliferation and differentiation in liver, gastroesophageal cancer and hematopoietic systems, and also to promote stemness of melanoma, glioma and colon cancers [36–41]. OPN overexpression correlates with the worst prognosis of human gastric and gastroesophageal carcinomas [41, 42].

OPN has been reported as a biomarker in ovarian cancers based on its detection in blood, primary and metastatic carcinomas [43–48]. It has been reported that acquired expression of OPN by HO-8910 cells greatly promotes soft agar colony formation and tumor growth in vivo [49]. Our study uncovers roles of OPN as a local niche factor in HGSC, and its effect on stem cell properties and chemo-resistance of ovarian cancer cells.

A key OPN receptor is CD44, which is a polymorphic transmembrane glycoprotein expressed in multiple cell types [50]. This cell surface receptor participates in a wide variety of cellular functions, such as cell growth, differentiation, apoptosis and motility [51, 52]. CD44 is expressed by Lgr5+ stem cells in the intestinal crypts [53] and stem/progenitor cells in gastric squamous-columnar junction[41]. It is also used as the marker for mesenchymal stem cells and hematopoietic stem cells [54, 55]. CD44 is expressed in KRT5+ human tubal epithelial with stem-like properties [27]. Thus, future studies should evaluate if OPN-CD44 signaling is a potential target for preventive or therapeutic intervention of cancers.

Recent developments in cancer therapy have targeted chemoresistant cancer propagating cells with a limited efficacy. Our studies point to the importance of non-CPCs in CPC maintenance. Compound targeting pathways that support stem-like (KRT5+ OPN-responsive CPC) and niche (KRT5- non-tumorigenic OPN-producing non-CPC) cell phenotypes may lead to maximizing effects of HGSC therapeutics. Furthermore, this approach may reduce therapeutic doses of highly toxic anticancer drugs.

## Methods

### Study groups and clinical data

Kaplan-Meier Plotter [56] was used as a survival analysis tool to evaluate *KRT5* mRNA expression in 470 patient samples obtained from The Cancer Genome Atlas (TCGA https://www.cancer.gov/ccg/ research/genome-sequencing/tcga) and other publicly available sources. KRT5 immunohistochemical staining was performed on tissue arrays prepared at the Institute of Pathology, Ludwig Maximilians University, Munich, Germany. Patients’ ages ranged from 37 to 100 years, with a mean age of 66 years. All patients were diagnosed with HGSC. For each case, a representative paraffin block containing invasive cancer tissue was selected for immunohistochemical analysis.

### Cell culture

Human ovarian cancer cell lines SKOV3 (ATCC, HTB-77), CAOV3 (ATCC, HTB-75), and CAOV4 (ATCC, HTB-76) were obtained from the American Type Culture Collection. The SKOV3 cells were cultured in McCoy’s 5A medium (Corning, 10-050-CV) supplemented with 10% Fetal Bovine Serum (FBS, Sigma, 12306C), 4 mM L-Glutamine (Corning, 25-005-Cl) and 100 IU ml^-1^/100 µg ml^-1^ penicillin/streptomycin (PS, Corning, 30-002-Cl).

CAOV3 cell line was maintained in Dulbecco’s Modified Eagle Medium (DMEM, Corning, 15-018-CV) supplemented with 10% FBS, 4 mM L-Glutamine and 100 IU ml^-1^/100 µg ml^-1^ PS. The culture conditions for SKOV3 and CAOV3 were 95% air, 5% carbon dioxide (CO_2_), at a temperature of 37°C. The COAV4 cell line was maintained in a Leibovitz’s L-15 (Corning, 10-045-CV) growth medium supplemented with 100 IU ml^-1^/100 µg ml^-1^ PS in 100% air, at a temperature of 37°C. HEK293T (Packaging cells, Human kidney, ATCC, CRL-3216) cells were cultured in DMEM, with 10% FBS, 1% non essential amino acids (NEAA, Thermo Fisher, 11140-050), 1 mM sodium pyruvate (Corning, 25-000-Cl), in 95% air, 5% CO2, at 37°C. Cell lines were authenticated in house and tested for *Mycoplasma* contamination every 6 months. Cell lines were used within 20 passages of thawing.

### Lentiviral packaging, transduction and shRNA mediated knockdown experiments

We performed lentiviral packaging and transduction following previously described methodology [57]. Briefly, for lentivirus packaging psPAX2 (Addgene, 12260), pMD2.G envelope plasmid (Addgene, 12259), and KRT5 promoter clone (GeneCopoeia, HRPM15909-LvPM02, mCherry) were employed. HEK293T cells were transfected with 10 µg KRT5 promoter clone, 7.5 µg psPAX2 and 2.5 µg pMD2.G using Lipofectamine 2000 (Thermo Fisher, 11668027) following manufacturer instructions. To differentiate between KRT5+ and KRT5- cell populations, another aliquot of transfection mixture containing ubiquitous promoter that drives GFP (hUbC, Addgene, 14883), was prepared in the same way. Each transfection mixture was added to a culture plate containing at least 3 x 10^5^ HEK293T cells. The culture plates were incubated for 12 hours, after which an additional 1 ml of antibiotic-free DMEM medium was added. Lentiviral supernatants were harvested after 48 hours and concentrated using centrifugal filters (Amicon, Ultra - 15, UFC903024). Freshly prepared lentiviral particles were used to transduce SKOV3 cells. Four days post transduction, viral media was replaced with SKOV3 growth medium containing 1 µg ml^-1^ puromycin to select for KRT5+ (mCherry^+^GFP^+^) and KRT5- (mCherry^-^GFP^+^) cell populations. Selected cells were cultivated for 2 passages, and either frozen in cryopreservation medium for storage at −80°C or immediately used for downstream applications. For gene silencing experiments, commercially available lentiviral particles with OPN (*Spp1*) shRNA (Santa Cruz Biotechnology, sc-36129-V) and scrambled control shRNA (Santa Cruz Biotechnology, sc-108080) were used to knockdown *Spp1* expression in SKOV3 cells and patient-derived cancer organoids.

### Fluorescence-activated cell sorting

For separation of KRT5+(mCherry+GFP+) and KRT5- (mCherry-GFP+) cell populations, the transduced SKOV3 cells were placed in Fluorescence-activated (FACS) cell sorting buffer (Phosphate buffered saline pH 7.4 (PBS), Corning 21-031-CV, + 1% bovine serum albumin, (BSA), Sigma A9647). Cell sorting and subsequent data analysis were performed on a FACS Aria II sorter that is equipped with the FACS DiVa software (BD Bioscience). Dead cells were excluded with Cytox Red (Thermo Fisher Scientific, S34859) and un-transduced SKOV3 cells served as controls. The brightest KRT5+ (mCherry+GFP+) and KRT5- (mCherry-GFP+) were identified and gated electronically based on their characteristic light-scattering properties on the mCherry and GFP channel emission pattern. The two populations were collected in two different tubes containing FACS cell sorting buffer and further stored at −80^°^C or immediately used for downstream applications. Fractions of L-KRT5mCherry (mCherry) and L-hUbC-GFP (GFP) DNA were detected by PCR with primers shown in Table S4.

### Single cell tracing

KRT5+ (mCherry^+^GFP^+^) and KRT5- (mCherry^-^GFP^+^) cell populations from SKOV3 cell line were suspended in different tubes containing culture medium and further counted to use them for single cell tracing experiment. Medium containing cells was diluted to contain a single cell per 0.5 ml culture medium and seeded in a 35 cm in diameter bottom glass dish. Cells were incubated at 37^°^C and 5% CO_2_. After attachment, the differentiation of a single cell was traced for 10 days using a fluorescent microscope that was equipped with mCherry and eGFP channels (Zeiss Axiovert 200 Live-Cell Incubation Fluorescence Microscope). The imaging data was processed using ImageJ and a movie was compiled for each KRT5+ and KRT5- cell.

### Cancer organoid culture

Primary ovarian tumors were acquired immediately during the surgery, and a part of the tumor was collected for organoid culture as described previously [58] or fixed immediately in 4% paraformaldehyde for later histological assessment. For organoid culture initiation the cryopreserved tissues were thawed in vials for 2–3 minutes in a 37°C water bath. Afterwards, in sterile conditions, washed three times in PBS, placed into 2 ml 2D media (Gibco Advanced DMEM/F12, Thermo Fisher Scientific, 12634028) supplemented with 5% FBS, 12 mM Hepes (Thermo Fisher Scientific, 15630080), 1% L-GlutaMax-I (Thermo Fisher Scientific, 35050079), 100 IU ml^-1^/100 µg ml^-1^ PS,, 10 µM Rho Kinase inhibitor (Y-27632, ROCKi, 1 mM, Millipore, 68800), 10 ng ml-1 epidermal growth factor human (EGF, 10 µg ml-1, Sigma, E9644) and then minced into 0.1 mm pieces using surgical blades.

The minced tissues were collected into 10 ml PBS, centrifugated at 300xg for 7 minutes at +4^°^C. The supernatant was discarded and the cell pellet suspended in 10 ml digestion buffer (0.5 mg/ml collagenase type I (Gibco 17100-017, in Advanced DMEM/F12, 12 mM Hepes, 3 mM CaCl_2_) followed by 45 minutes incubation at 37^°^C in a water bath. After the incubation 30 ml of 2D medium was added to the 10 ml mixture, and the solution was passed through 100 µm (Falcon; #352360), 70 µm (Falcon; #352350), and 40 µm cell strainers (Falcon; #352340). Next, the suspension was centrifuged at 300xg for 7 minutes at +4^°^C. The cell pellet was suspended in 2D media, counted and adjusted to 5 × 10^4^ cells per rim assay. Matrigel (Corning; 356231) was added to a final concentration of Matrigel at 70%. Rim assays were seeded in rims of 24 well plates and cultures performed as previously reported [58].

FACS sorted SKOV3 cell line-derived KRT5+ and KRT5- cells were suspended in 2D media so that the suspension contains 5 × 10^4^ cells. Subsequently, organoid cultures were performed as previously described [58].

### Cell viability assays

Cell viability after administration of either Cisplatin (Cis-Diammineplatinum(II) dichloride; Pt(NH_3_)_2_Cl_2_, Sigma, P4394) or Doxorubicin (Sigma, 44583) was established by manual count and 3-(4,5)-Dimethylthiahiazo(-z-y1)-3,5-di-phenytetrazoliumromide (MTT) assays. For manual count cells were seeded into 6-well plates at 5.0 x 10^4^ cells. The MTT assay was carried out as per the manufacturer’s instructions (Abcam, #ab211091). Briefly, cells were seeded at a density of 5.0 x 10^4^ cells/well in 96 well plates. The media was discarded from cell cultures and 50 µl of serum-free media and 50 µl of MTT solution were added to each well. The plates were incubated at 37^°^C for 3 hours. After incubation, 150 µl of MTT solvent was added to each well and incubated on orbital shaker for 15 minutes and the absorbance was read at OD = 590 nm. All experiments had a minimum of three replicates.

### Immunohistochemistry and quantitative image analysis

Immunohistochemical analysis of paraformaldehyde-fixed paraffin embedded tissues was performed using a modified ABC technique [59–61]. Briefly, the ovarian cancer patient derived tumors and mouse xenografts were excised and fixed with 4% paraformaldehyde and incubated overnight at 4^°^C. Tissues were dehydrated and embedded using paraffin and sectioned 5 µm thick and subjected to IHC staining. The slides were deparaffinized in xylene and rehydrated over a graded ethanol series, and antigen retrieval was performed using citrate buffer at pH 6.0. All primary and secondary antibodies for immunohistochemistry are shown in Table S5. The VECTASTAIN ABC reagent (Vector Laboratories, PK-6100) and 3,3-Diamino benzidine (DAB, Sigma 1029240001) were used for signal detection. Sections with no primary antibody served as negative controls. Tissues sections with immunoperoxidase staining were scanned and digitized using ScanScope CS2 (Aperio Technologies) with a 40x objective. Immunofluorescent stained sections were mounted in ProLong Diamond Antifade Mountant with DAPI reagent (Thermo Fisher, P36962)) and sealed using nail polish and further scanned using ScanScope FL (Aperio Technologies) with a 20x objective. Further, ImageJ software (National Institute of Health, NIH) was used for quantitative analysis of immunohistochemistry and immunofluorescence images as previously described [59–61].

For immunocytochemistry in cell culture, cells were grown on coverslips and fixed using 4% paraformaldehyde in PBS pH 7.4 for 10 min at room temperature. Permeabilization was performed using PBS containing 0.1% Triton X-100 for 10 min and washed in PBS three times for 5 min. The permeabilized cells were incubated with 1% BSA, 22.52 mg/mL glycine in PBST (PBS+ 0.1% Tween 20) for 30 min to block nonspecific proteins. Further cells were incubated with primary antibody overnight at 4^°^C. Primary and secondary antibodies used to stain cells are listed in Table S5. The cells were washed three times with PBS for 5 min and incubated with secondary antibody for 1 hour at room temperature in the dark. Stained cells were mounted in the same way as immunofluorescent stained tissue sections. Confocal images were acquired using a Zeiss LSM 710 confocal microscope through the Cornell University Biotechnology Resource Center. The image data was merged and displayed with the ZEN software (Zeiss).

### Quantitative real-time PCR

RNA was extracted using TRIzol reagent according to manufacturer’s instructions (Thermo Fisher, 15596026). cDNA was produced using the SuperScript III First-Strand Synthesis kit (Thermo Fisher, 18080400). Real-time PCR was performed using PerfeCTa SYBR Green Super Mix Reagent (Quanta Biosc., 95072-250) on C1000 Touch Thermal Cycler PCR machine (Bio-Rad).

### Tumorigenicity assays

KRT5+ and KRT5- cancer cells FACS-isolated from transduced SKOV3 cell line were suspended in PBS and injected subcutaneously (5 x 10^5^, 5 x 10^4^, and 5 x 10^3^ cells in 250 µl) into either side of flanks of 6-8-week-old *NOD.Cg-Prkdc^scid^Il2rg^tm1Wjl^*/*SzJ4* female mice. Control mice received only PBS. The size of the subcutaneous mass was measured using scientific caliper every alternate day starting from day 20 after transplantation until day 30. The tumor volume was calculated using VT = 1/2(L x W x H) and tumor incidence was also noted. Mice were euthanized and subjected to necropsy as soon as they showed signs of sickness such as abdominal distension, subcutaneous masses approximately 1 cm diameter on either side of flanks or moribund behavior.

### RNA sequencing

Total RNA was isolated using TRIzol reagent according to manufacture’s instructions. RNA integrity and concentrations were determined using Nanodrop and only the purified RNA samples were used for further analysis. ERCC Spike in in Mix 1 (Thermo Fisher 4456740) was added to samples prior to library creation according to manufacturer’s recommendation. 3’ mRNA-seq libraries were created using QuantSeq 3’ mRNA-Seq library prep kit (New England Biolabs) according to the manufacturer’s instructions using 10 ng total RNA. The sequencing files were inspected for quality control analysis using FASTQC [62]. Reads were then aligned to the GRCh38 human reference genome using the STAR two-pass method. R (https://www.r-project.org/) was used for all analysis and visualization of aligned data. The differential gene-expression analysis was performed using the R package DEseq2 (V 1.6.3). A false discovery rate of ≤0.05 was then used to select for differentially expressed genes between sample group comparisons. Heat maps were generated using normalized average gene-count values within the groups. Only the genes that were significantly differed (p < 0.05) between KRT5+ and KRT5- groups were taken into consideration for further analysis. Further, the Database for Annotation, Visualization and Integrated Discovery (DAVID) (v6.8) gene functional annotation and classification tool [63, 64] was used to annotate the list of differentially expressed genes with respective GO terms and to perform GO enrichment analysis and gene functional clustering for biological functions including significant pathways and gene ontology terms. Statistically significant signaling pathways were then identified by using a P value cut off of 0.05.

### Single-cell RNA sequencing analysis

A publically available dataset for 29 HGSC cases [GSE180661][65] were downloaded and aligned to the GRCh38 human reference genome using cellranger (v7.1.0, 10x Genomics). All further preprocessing and data analysis was conducted in R (v4.2.0). The count matrices were all modified with the standard SoupX (v.1.6.1) pipeline to remove ambient RNA signal (https://github.com/constantAmateur/SoupX). The corrected matrices were then processed in Seurat (v.4.3.0) (https://github.com/satijalab/seurat) to subset epithelial cells and quantify based on *KRT5* and *SPP1* expression.

### Statistical analysis

Statistical analysis was performed on GraphPad Prism 10.6.1. All in vitro experiments were performed in biological triplicates unless otherwise stated. Two-tailed Mann-Whitney test and unpaired t test with Welch’s correction were used to compare two groups. ANOVA was conducted for multiple comparisons followed by a post hoc Tukey test to identify differences within groups. All data are shown as the mean with SDs from at least three independent experiments. Probabilities of *P* > 0.05 were considered as not significant, and significance was established as *, *P* < 0.05; **, *P* < 0.01; ***, *P* < 0.001; ****, *P* < 0.0001. Survival analysis was conducted using the Kaplan–Meier method. *P* < 0.05 indicated significant difference.

## Declarations

### Ethics Statement

De-identified tissue microarrays of HGSC were prepared at the Institute of Pathology, Ludwig Maximilians University, Munich, Germany for studies approved by the Ethics Committee of the Ludwig-Maximilian-University, Munich, Germany. De-identified clinical samples used for HGSC organoids were obtained from Weill Cornell Medicine (Cornell IRB0005546) and from University of Kansas Cancer Center’s Biospecimen Repository Core Facility (BRCF) at the KU Medical Center together with relevant clinical information. HGSC tissue specimens were obtained from women enrolled under the repository’s IRB approved protocol (HSC #5929) and in accordance with the U.S. Common Rule. All patients provided written informed consent in compliance with the BRCF IRB protocol. Samples were de-identified using OpenSpecimen prior to analysis. Unless otherwise indicated, all sample handling and experimental procedures were conducted in a biosafety level 2 (BSL-2) cabinet.

### Availability of Data and Material

The bulk RNA-seq data reported in this paper is deposited in Gene Expression Omnibus (GEO); accession number (GSE316118). The Source Data provides data for all results requiring quantification. Any additional data supporting the findings of this study are available from the corresponding author upon reasonable request.

### Author contributions

Conceptualization: M.B., A.F.N., A.Y.N.; Methodology: A.F.N., A.Y.N.; Software: C.Q.R., I.M.R.; Validation: M.B., D.C., A.F.N., A.Y.N.; Formal Analysis: M.B., C.Q.R, I.M.R., A.F.N., A.Y.N.; Investigation: M.B., C.Q.R, N.B., I.M.R., A.F.N., A.Y.N.; Resources: E.S.,A.K.G.,D.M.,L.H.E.; Data Curation: M.B., A.F.N., A.Y.N.; Writing – Original Draft: M.B., A.F.N., A.Y.N; Writing – Review & Editing: M.B., C.Q.R, A.F.N., A.Y.N.; Visualization: M.B., C.Q.R, A.F.N., A.Y.N.; Supervision: A.F.N., A.Y.N.; Project Administration: A.F.N., A.Y.N.; Funding Acquisition: M.B., C.Q.R., I. M. R., A.Y.N.

## Acknowledgements

We thank Peter A. Schweitzer, Director of the Cornell Genomics Facility for his RNA sequencing, Paris Zhang for her help with a pilot chemoresistance project, and all members of the Nikitin Lab for their advice and support.

