## Supplementary Materials for "Keratin 5 marks cancer-propagating cells sustained by an osteopontin-producing niche in high-grade serous ovarian carcinoma"

### Table S1. Frequency of KRT5+ cells in HGSC cell lines according to immunostaining.

| Cell Line | N | Mean ± SD (%) |
|---|---|---|
| SKOV3 | 4 | 7.91 ± 2.32 |
| CAOV3 | 6 | 2.43 ± 2.37 |
| CAOV4 | 4 | 4.97 ± 1.47 |

### Table S2. Frequency of KRT5+ cells in primary HGSC according to scRNA seq datasets.

| HGSC group | N | Mean ± SD (%) |
|---|---|---|
| HGSC (>0%) | 29 | 10.87 ± 13.94 |
| HGSC (>5%) | 14 | 20.15 ± 15.42 |

### Table S3. Tumor incidence.

| Injected Cells | Injected mice/Mice with tumors | | |
|---|---|---|---|
| | $1 \times 10^3$ | $1 \times 10^4$ | $1 \times 10^5$ |
| KRT5+ | 4/4 | 5/5 | 6/6 |
| KRT5- | 0/5 | 0/5 | 0/6 |

### Table S4. Genotyping primers.

| Primer name | Primer sequence 5'-3' | PCR product size (bp) |
|---|---|---|
| GFP GT 346n-Forward | CCTCGTGACCACCCTGACCTACGGC | 350 |
| GFP GT 346n-Reverse | GCCGTCCTCGATGTTGTGGCGGATC | |
| mCherry_9703_Forward | AGGACGGCGAGTTCATCTAC | 288 |
| mCherry_9704_Reverse | TGGTGTAGTCCTCGTTGTGG | |

### Table S5. Antibodies used for immunostaining.

| Antibodies | Source | Identifier |
|---|---|---|
| Mouse monoclonal [XM26] to Cytokeratin 5 | Abcam | ab17130data |
| Anti-SPP1 | Sigma | HPA027541 |
| Mouse monoclonal to mCherry | Abcam | ab125096 |
| Alexa Fluor 488 Donkey anti-Mouse | Life Technologies | A-21202 |
| Alexa Fluor 488 Donkey anti-Mouse | Life Technologies | A-21207 |
| Alexa Fluor 594 Donkey anti-Rabbit | Life Technologies | A-21207 |

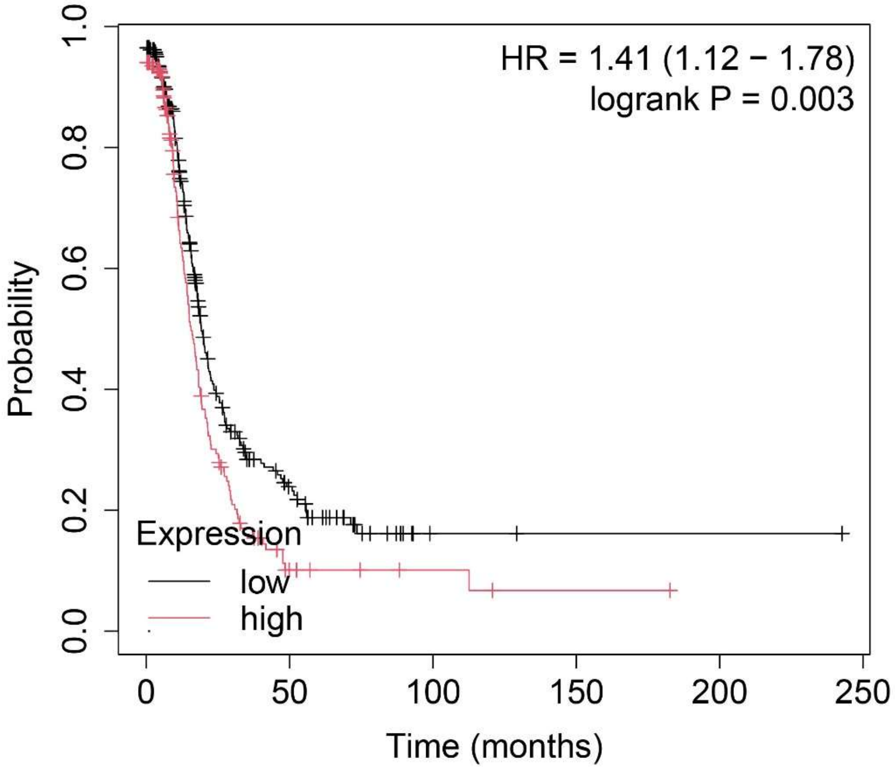


**Figure S1. Overall survival of 470 patients with serous ovarian carcinoma with *TP53* mutation according to KRT5 mRNA expression.** KM Plotter analysis.

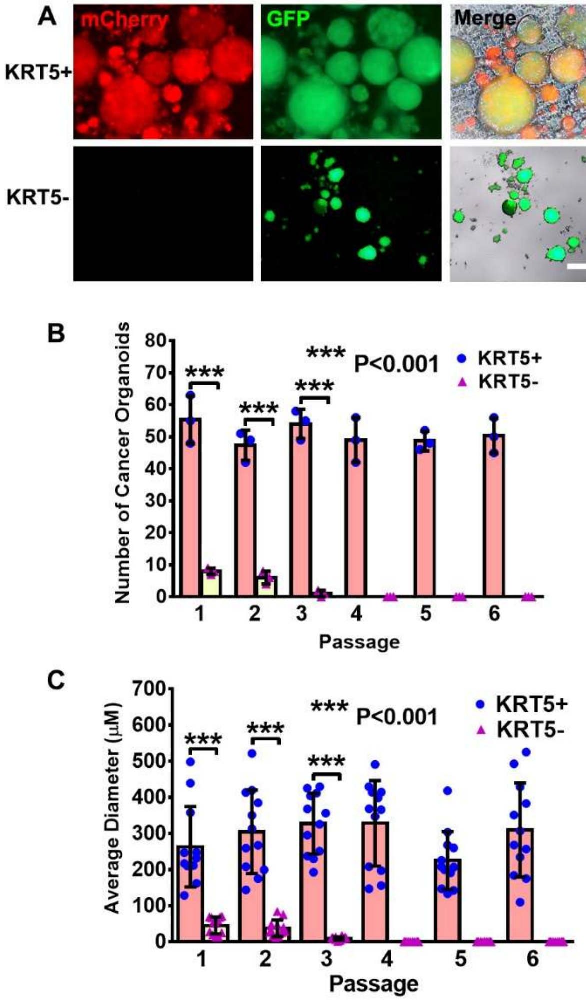


**Figure S2. Characterization of KRT5+ and KRT5- cell derived cancer organoids in ultra-low attachment plates. A.** Representative images of KRT5+ and KRT5- cell derived cancer organoids. Bar, 100 µm for all images. **B.** Quantification of number of KRT5+ and KRT5- cancer organoids in for six consecutive passages. **C.** Average diameter of KRT5+ and KRT5- cancer organoids. in Matrigel for six consecutive passages. All error bars denote s.d.
